# A telosma mosaic virus-based vector for foreign gene expression and virus-induced gene silencing in the perennial woody vine, passion fruit (*Passiflora edulis)*

**DOI:** 10.1101/2024.08.13.607796

**Authors:** Xiaoqing Wang, Li Qin, Wentao Shen, Wenping Qiu, Hongguang Cui, Zhaoji Dai

## Abstract

Passion fruit (*Passiflora edulis*) is a perennial, woody, tropical vine crop. It produces edible round to oval fruit that has been increasingly favored for its unique aroma and taste, and richness in antioxidants, vitamins and minerals. However, the functional genomic study of passion fruit lags far behind due to a lack of simple and efficient genetic tools. Here, we report the development of virus-mediated protein overexpression (VOX) and virus-induced gene silencing (VIGS) vector based on telosma mosaic virus (TelMV), an emerging potyvirus infecting passion fruit plants worldwide. This vector, designated pTelMV-GW, incorporates the Gateway-compatible recombination sites for rapid gene cloning. We show that this vector allows for the systemic stable expression of two heterologous proteins, green fluorescent protein (GFP) and bacterial phytoene synthase (crtB) in passion fruit plants, and pTelMV-GW containing different fragments of *GFP* can also induce systemic gene silencing on the GFP-transgenic *N. benthamiana* plants. Moreover, we demonstrated that in passion fruit plants, this vector can trigger gene silencing of endogenous *phytoene desaturase* (*PDS*) to a limited extent. Furthermore, we upgraded the vector by using a mild TelMV strain that does not induce noticeable symptoms in plants. We show that the upgraded vector (pTelMV-R181K-GW) containing *PDS* or *ChlI* fragments induces the robust silencing of the corresponding endogenous gene in passion fruit plants. Together, we reported the first development of VIGS and VOX vectors in passion fruit plants, as the first step in our endeavor to discover horticulturally important genes for improving passion fruit production and quality.

## Introduction

Passion fruit (*Passiflora edulis*) is a perennial evergreen woody vine grown in sub-tropical and tropical areas. It belongs to the *Passiflora* genera in the family of *Passifloraceae* (Xia et al., 2021; Wang et al., 2023). Over the years passion fruits have become the new favorites in both drinks and foods as it contains multiple vitamins and antioxidants that benefit human health. It was not until 2021 that a chromosome-scale genome assembly of passion fruit was just reported (Xia et al., 2021) and the functional genomic study of passion fruit plants has not been fully initiated. The functions and regulations of genes engaged in the biosynthesis of the health-promoting compounds in passion fruit remain largely unknown. This is mainly due to the lack of simple and efficient tools for functional genomic analysis in passion fruit.

Plant viruses have been engineered as vectors for virus-mediated protein overexpression (VOX) and virus-induced gene silencing (VIGS), as they comprise relatively small genomes, are easy to manipulate, replicate rapidly and spread throughout the whole plant (Constantin et al., 2004; Cui et al., 2017). As a reverse genetic tool, VOX and VIGS are powerful alternative approaches for determining gene function in plant species that are not amenable to the classic genetic transformation methods, such as *Agrobacterium*-mediated transformation (Constantin et al., 2004; Hileman et al., 2005). In addition, VOX and VIGS possess the great advantage of being fast that the time from cloning the gene of interest (GOI) to the phenotype analysis is much shorter compared to stable plant transformation. Despite the fact that the *Agrobacterium*-mediated stable transformation and regeneration of passion fruit have been established (Trevisan et al., 2006; Correa et al., 2015; Rizwan et al., 2021), the process is time-consuming, laborious and inconvenient for high-throughput characterization of passion fruit genes. Unfortunately, VOX and VIGS vectors have not been created in passion fruit plants.

Telosma mosaic virus (TelMV) is an emerging virus infecting passion fruit plants in multiple plantations in China, Brazil, Vietnam and Japan (Gou et al., 2023; Wang et al., 2024). Since the discovery of TelMV in telsoma (*Telosma cordata*, *Asclepiadaceae*) in 2008, the host range of TelMV has been continuously expanding to multiple species, including patchouli, pigweed, kidney bean, quinoa, Emperor’s candlesticks, and tobacco (Zhang et al., 2024). TelMV belongs to the genus *Potyvirus* in the family *Potyviridae*. Potyvirus represents the largest group of plant RNA viruses and comprises a positive-sense single-stranded RNA genome encapsidated by viral coat protein. The potyvirus genome encodes a large polyprotein that is further cleaved into ten mature viral proteins by its viral proteases. In addition, potyvirus also encodes a short ORF (pretty interesting *Potyviridae* ORF, PIPO) resulting from RNA polymerase slippage during viral replication (Revers and García, 2015; Cui and Wang, 2019; Dai et al., 2020; Yang et al., 2021; Wu et al., 2024). Although potyviruses have been broadly engineered into protein expression vectors, such as tobacco etch virus (TEV), potato virus A (PVA), turnip mosaic virus (TuMV), plum pox virus (PPV), sugarcane mosaic virus (SCMV), wheat streak mosaic virus (WSMV), tobacco vein banding mosaic virus (TVBMV), zinnia mild mottle virus (ZiMMV), maize dwarf mosaic virus (MDMV) and zucchini yellow mosaic virus (ZYMV) (Xie et al., 2021; Yang et al., 2024), very few potyviruses have been reported for VIGS vector (only four cases: papaya leaf distortion mosaic virus, PLDMV; MDMV; watermelon mosaic virus, WMV and SCMV) (Tuo et al., 2021; Xie et al., 2021; Houhou et al., 2021; Chung et al., 2022). This is mainly due to two factors, one is that potyvirus encodes a strong RNA silencing suppressor HC-Pro that inhibits the host RNA-based antiviral response (Valli et al., 2018). Secondly, potyvirus employs polyprotein processing as its gene expression strategy. Pioneering studies of potyviruses have demonstrated the manipulation of the potyviral genome is mainly restricted to the sites for inserting a non-viral fragment in the P1/HC-Pro and NIb/CP junctions (Xie et al., 2021; Houhou et al., 2021). Our lab has constructed the first passion fruit-infecting TelMV infectious clone pPasfru and it can tolerate foreign insertion at the NIb/CP junction (Gou et al., 2023).

Typically, VOX and VIGS involve the cloning of the full-length or fragment of GOI into the viral vector, viral infection of the plant hosts, and protein expression/silencing of the target genes by the RNA-mediated antiviral defense mechanism in plants (Rossner et al., 2022). Since the first report of VIGS in 1995 (Kumagai et al., 1995), VIGS has been proven to be an efficient tool for gene function analysis in diverse plant species, including tobacco, soybean, wheat, tomato, Arabidopsis and petunia (Dommes et al., 2019; Rossner et al., 2022). To facilitate convenient and rapid gene cloning, Gateway-based strategy has been incorporated into VIGS vectors. In this system, the Gateway recombination cassette *att*R1-Cm^R^-*ccd*B-*att*R2 is inserted in the viral vector, thus allowing the simple, fast and high-throughput cloning of GOI. Multiple viral vectors compatible with the Gateway cloning technology have been reported for VOX and VIGS in plants, including tobacco rattle virus (TRV), potato virus X (PVX), tobacco mosaic virus (TMV), and cymbidium mosaic virus (CymMV) (Liu et al., 2002; Lacorte et al., 2010; Lu et al., 2012). The most broadly-used VIGS vector is the TRV-based vector. For instance, the Gateway-compatible TRV VIGS vector was successfully applied to silence endogenous genes in tomatoes (Liu et al., 2002).

This study aims to develop TelMV-based VOX and VIGS systems in passion fruit plants. Here, we engineered a Gateway-compatible TelMV VOX vector that can systemically express heterologous proteins, green fluorescent protein (GFP), and bacterial phytoene synthase (crtB) in passion fruit. In addition, we developed the TelMV-based VIGS vector that can efficiently silence the endogenous phytoene desaturase (*PePDS*) gene and magnesium chelatase subunit I (*PeChlI*) gene in passion fruit plants. The TelMV-derived VOX and VIGS tools described here constitute ideal tools for functional genetic analysis in passion fruit plants, especially the genes in the biosynthesis of the health-promoting compounds in passion fruits.

## Results

### Development of telosma mosaic virus-based vector compatible with Gateway cloning technology

To develop a VIGS vector for passion fruit plants, we first tested whether the most commonly used TRV vector could be utilized for VIGS. Unfortunately, TRV could not infect passion fruit plants using either the agroinfiltration or rub-inoculation method in our current experimental setup (data not shown). We then decided to use telosma mosaic virus (TelMV), a natural passion fruit-infecting potyvirus, and engineered it into a viral vector that is compatible with the Gateway cloning technology for rapid gene cloning.

To this end, the Gateway recombination frame (GW Frame) comprising the classic *att*R1-*Cm^R^-ccd*B -*att*R2 recombination cassette flanked by two NIa-Pro cleavage sites was placed between the NIb/CP junction (replacing GFP cistron) in the backbone of TelMV infectious clone pPasFru-G (pTelMV-GFP) (Gou et al., 2023) (Figure 1A), allowing the release of the foreign proteins from the viral polyprotein. The resulting construct was designated pTelMV-GW (Figure 1B). We included silent mutations in the second NIa-Pro cleavage site to avoid long sequence repetitions, minimizing the risk of homologous recombination during virus replication. Lastly, we verified the introduced features of the new pTelMV-GW vector by PCR, double digestion and Sanger sequencing (data not shown) to ensure that GOI can be cloned into the vector through one-step LR reaction (Figure 1C).

**Fig 1.**
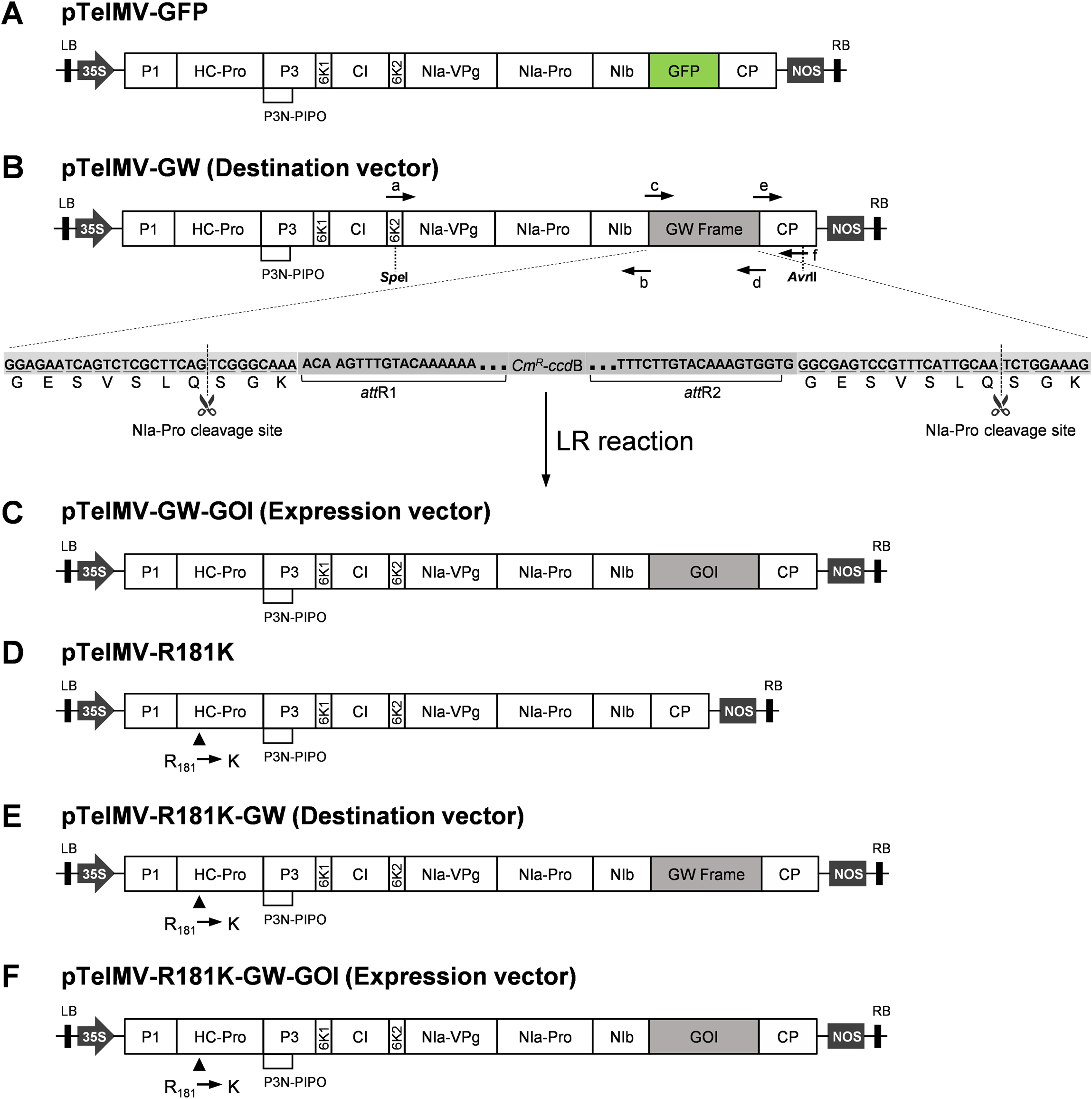
Schematic representation of the Gateway-compatible TelMV-based vectors. **A.** Schematic representation of telosma mosaic virus (TelMV) infectious clones, pTelMV-GFP. LB, RB: left and right border of T-DNA; GFP: green fluorescent protein; 35S: 35S promoter of cauliflower mosaic virus; Nos: nopaline synthase terminator. **B.** Schematic representation of the Gateway-compatible TelMV vector, pTelMV-GW. The five letters (a, b, c, d, e, and f) represent the primers used for TelMV-GW cloning by overlapping PCR. The Gateway recombination frame was inserted between the viral nuclear inclusion protein b (NIb) and coat protein (CP) cistrons. The magnification shows the Gateway recombination frame (GW Frame) sequence that contains the last seven amino acid resides of NIb cistron (GESVSLQ), the first three amino acid resides of CP cistron (SGK) and the Gateway recombination cassette *att*R1-Cm^R^-*ccd*B-*att*R2. *att*R1, *att*R2: Gateway recombination sites; Cm^R^, *ccd*B, selection markers for Gateway cloning. **C.** Schematic representation of the Gateway-compatible TelMV-based expression vector, pTelMV-GW-GOI. GOI: gene of interest. **D.** Schematic representation of TelMV-R181K infectious clone (pTelMV-R181K), which harbors a point mutation (R to K) in the position of 181 of HC-Pro cistron. **E.** Schematic representation of Gateway-compatible TelMV-R181K-based vector, pTelMV-R181K-GW. **F.** Schematic representation of the Gateway-compatible TelMV-R181K-based expression vector, pTelMV-R181K-GW-GOI. GOI: gene of interest.

### Stable systemic expression of GFP using pTelMV-GW in passion fruit plants

To determine whether pTelMV-GW could systemically infect plants and express foreign protein upon insertion of GOI, the green fluorescent protein (GFP)-encoding gene *GFP* was cloned into the entry clone pDONR221, and subsequently inserted into pTelMV-GW upon LR recombination reaction. The resulting construct, pTelMV-GW-GFP (Figure 2A), was transformed into *Agrobacterium tumefaciens* and agro-infiltrated into the three-week-old *N. benthamiana* plant leaves. At 7 days post inoculation (dpi), green fluorescence was first observed in the systemic leaves under UV light, and the fluorescence became more intense at 9 dpi (Figure 2B), as in the plants inoculated with the previously constructed GFP-tagged TelMV infectious clone (pTelMV-GFP) (Gou et al., 2023) (Figure 2B). In contrast, no green fluorescence was detected in *N. benthamiana* plants infiltrated with buffer (Mock). Furthermore, the presence of GFP transcripts and protein was validated with RT-PCR and Western blot analyses, respectively, in the systemic leaves infiltrated with pTelMV-GW-GFP (Figure 2C and 2D, left panels). These data demonstrated that a gene of interest could be systemically expressed in *N. benthamiana* plants using the Gateway-compatible viral vector, pTelMV-GW.

**Fig 2.**
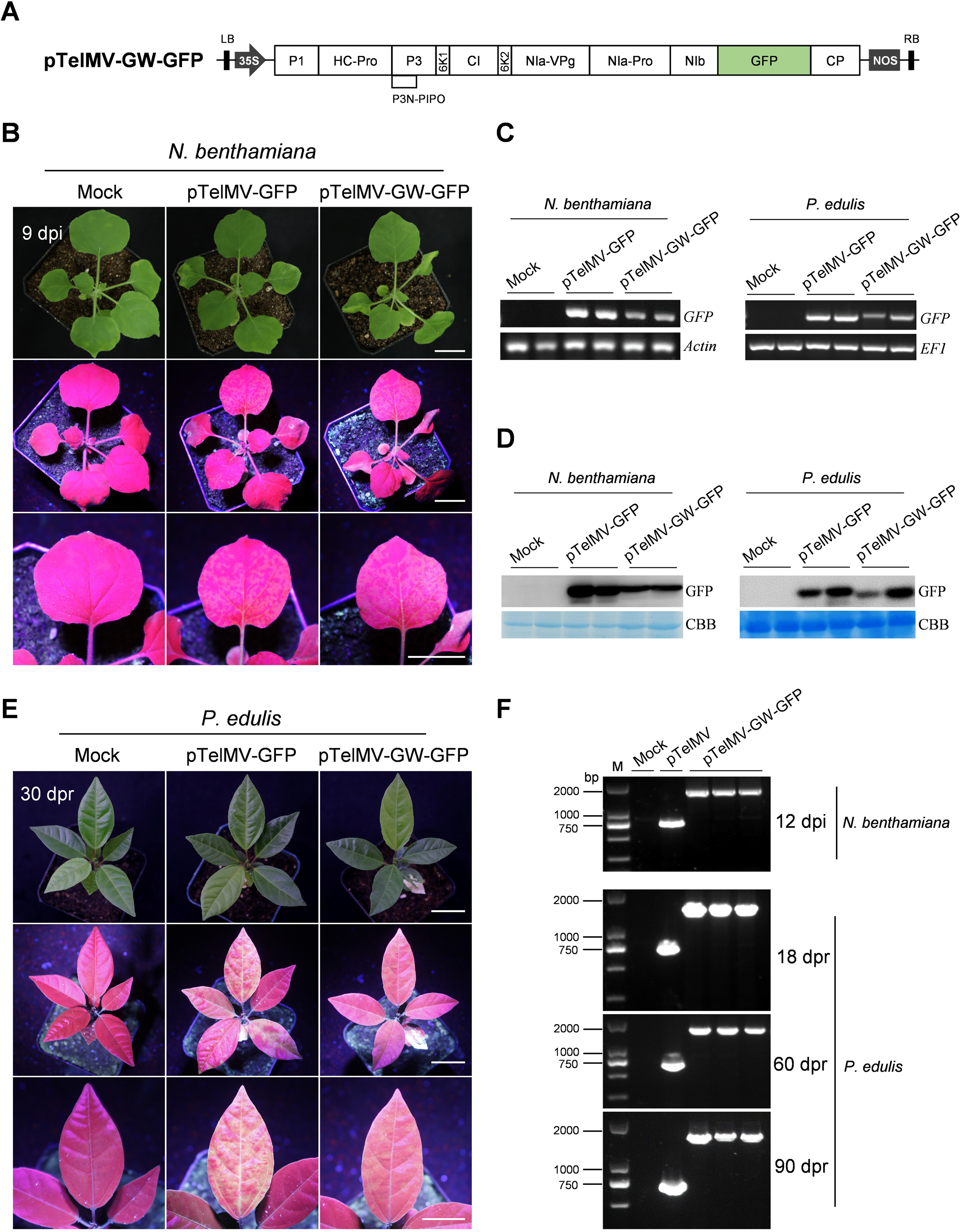
pTelMV-GW as a gene expression vector in *N. benthamiana* and *P. edulis* plants. **A.** Schematic representation of pTelMV-GW-GFP vector for GFP expression. GFP: green fluorescent protein. **B.** *N. benthamiana* plants inoculated by buffer (Mock), positive control (pTelMV-GFP) and pTelMV-GW-GFP, respectively under UV light at 9 dpi. Bar, 2 cm. **C.** Detection of GFP transcripts in both *N. benthamiana and P. edulis* inoculated with pTelMV-GW-GFP by RT-PCR analyses. The *N. benthamiana Actin* gene and passion fruit *EF1* gene serve as the internal controls. **D.** Detection of GFP protein in both *N. benthamiana and P. edulis* inoculated with pTelMV-GW-GFP by Western blot analyses. The Coomassie brilliant blue (CBB)-stained Rubisco large subunit (RbcL) serves as a loading control. **E.** *P. edulis* plants inoculated by buffer (Mock), positive control (pTelMV-GFP) and pTelMV-GW-GFP, respectively under UV light at 30 dpr. Note that clear green fluorescence was observed on the newly emerged leaf of *P. edulis* inoculated with pTelMV-GW-GFP upon UV illumination. **F.** RT-PCR analysis of RNA samples extracted from pTelMV-GW-GFP-infected plants at various days. M: DNA ladder. pTelMV plasmid DNA serves as the wild-type virus control.

Next, the sap prepared from pTelMV-GW-GFP-agroinfiltrated *N. benthamiana* leaves harvested at 5 dpi served as a virus inoculum for the rub-inoculation of *P. edulis* plants (passion fruit). The green fluorescence was first observed at about 7-12 days post-rub-inoculation (dpr), and strong green fluorescence was observed at 30 dpr (Figure 2E). Intense fluorescence was also exhibited in the systemic leaf of passion fruit plants inoculated by the positive control (pTelMV-GFP), but not the Mock plants (Figure 2E), suggesting pTelMV-GW-GFP can systemically infect the passion fruit plants. Moreover, RT-PCR and Western blot analyses confirmed the presence of mRNA and protein of GFP, respectively, in the upper leaf of passion fruit plants inoculated by pTelMV-GW-GFP (Figure 2C and 2D, right panels). These results demonstrated that pTelMV-GW can serve as an excellent expression vector in passion fruit plants.

To assess the stability of the inserted foreign gene overtime during virus multiplication, we carried out RT-PCR analyses on total RNA extracted from the systemic leaves both in *N. benthamiana* and passion fruit plants using appropriate primers flanking the insertion sites at 12 dpi and 18 dpr, respectively. The Mock plant and the pTelMV plasmid DNA (producing a 777-bp PCR fragment) served as the negative and positive controls (Figure 2F). A larger single band containing the intact GFP-coding sequencing (1575 bp) could be detected (Figure 2F), but not the deletion derivatives, suggesting the TelMV vector retained the GFP sequence upon viral systemic infection. In addition, no truncated variant progeny was detected at extended days, 60 dpr and 90 dpr, respectively by the RT-PCR analysis of samples harvested from the upper leaf of passion fruit plants (Figure 2F). These results showed that pTelMV-GW-GFP is stable during virus replication in plants up to 90 dpr.

Therefore, pTelMV-GW can be used for stable systemic expression of GOI in *N. benthamiana* and passion fruit plants.

### Overexpression of a heterologous protein CrtB from TelMV VOX vector led to yellowing of passion fruit leaf

To further investigate the capability of pTelMV-GW for expressing the functional protein in passion fruit plants, a Myc-tagged bacterial phytoene synthase gene (*crtB*) was cloned into the pTelMV-GW vector through the LR reaction. The resulting construct, pTelMV-GW-^myc^CrtB (Figure 3A), was transformed into *A. tumefaciens* and agroinfiltrated into *N. benthamiana* leaves for the subsequent rub-inoculation of passion fruit plants. The bacterial *crtB* gene encodes a phytoene synthase that catalyzes the condensation of two molecules of geranylgeranyl diphosphate (GGPP), thereby initiating the carotenoid biosynthesis pathway (Tuo et al., 2023). The overexpression of crtB gene in plants using viral vectors often resulted in the overaccumulation of carotenoids and the characteristic yellow phenotype of *crtB*-expressing leaf tissues (Llorente et al., 2020). As expected, pTelMV-GW-^myc^CrtB resulted in the yellowing of *P. edulis* leaves as early as 20 dpr and developed an intense yellowing phenotype at 30 dpr (Figure 3B). Meanwhile, the systemic expression of crtB was further validated by the presence of mRNA and protein using RT-PCR and Western blot analysis, respectively (Figure 3C).

**Fig 3.**
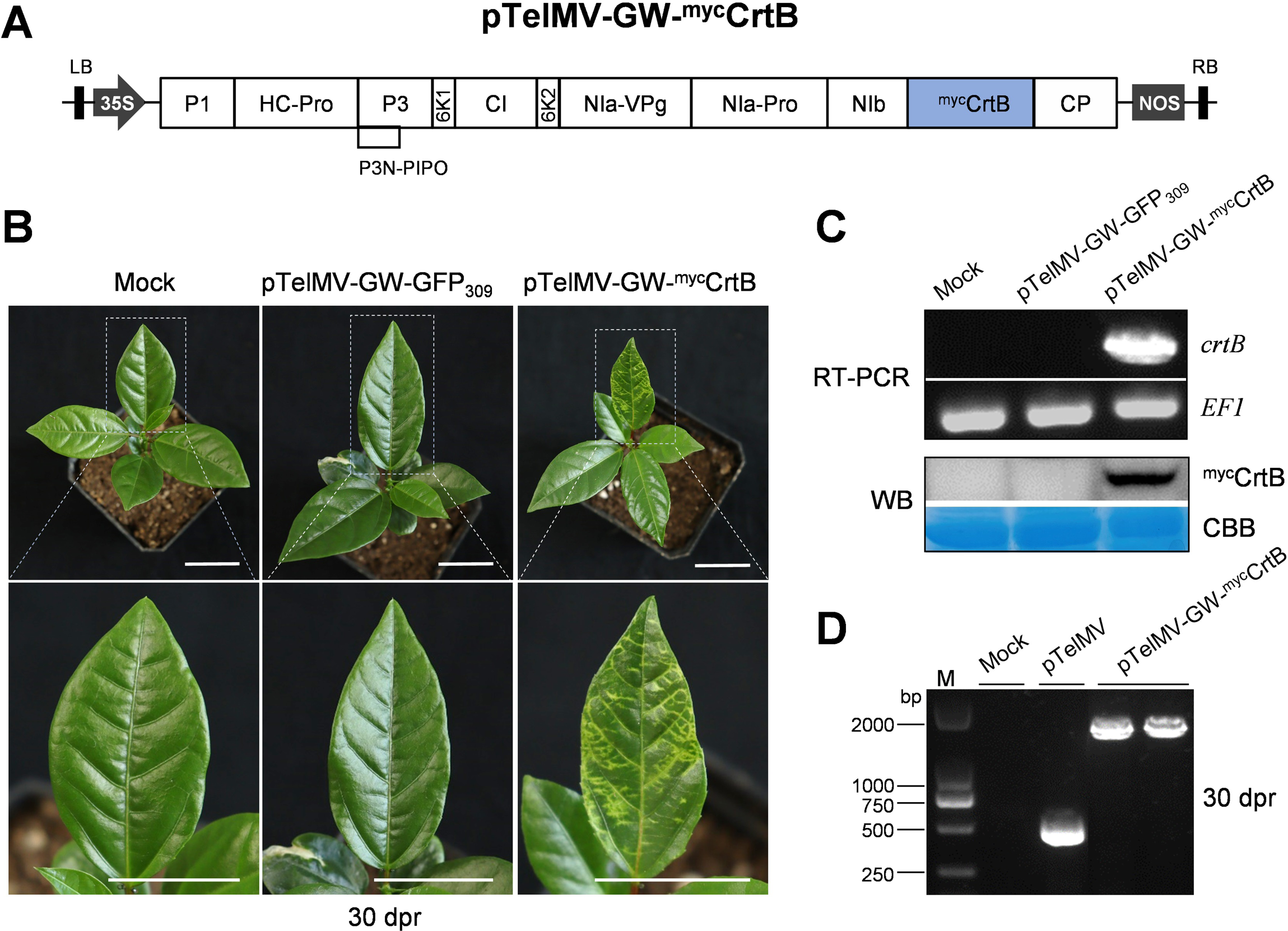
Expression of heterologous protein CrtB in passion fruit plants. **A.** Schematic representation of pTelMV-GW-^myc^CrtB vector for Myc-tagged CrtB protein production. *crtB*: bacterial phytoene synthase gene. **B.** The passion fruit plants inoculated by buffer (Mock), virus control (pTelMV-GFP_309_) and pTelMV-GW-^myc^CrtB, respectively, at 30 dpr. Bar, 2 cm. **C.** RT-PCR and Western blot analyses on the samples from passion fruit plant infected by pTelMV-GW-^myc^CrtB. The passion fruit *EF1* gene serves as an internal control. The Coomassie brilliant blue (CBB)-stained Rubisco large subunit (RbcL) serves as a loading control. **D.** RT-PCR analysis of RNA samples extracted from pTelMV-GW-^myc^CrtB -infected passion fruit plants at 30 dpr. M: DNA ladder. pTelMV plasmid DNA serves as the wild-type virus control.

The stability of crtB insertion was evaluated by the RT-PCR analysis of the systemic leaving exhibiting characteristic yellow phenotype using a set of primers flanking the crtB insertion. A single bright band containing crtB fragment was detected, but not the truncated versions in the two independent experiments at 30 dpr (Figure 3D). These results revealed that the crtB insertion is stable in the TelMV genome during virus replication in passion fruit plants.

These data further strengthened that a GOI can be systemically expressed by TelMV VOX vector for analyzing its function in the compound biosynthesis pathway in passion fruit plants.

### Visualization of VIGS in the 16c *N. benthamiana* plants

To examine the potentiality of the Gateway-compatible TelMV vector (pTelMV-GW) for VIGS, GFP-transgenic *N. benthamiana* plants (16c) were used for gene silencing analysis. The full length of GFP was divided into two parts, 309 bp and 405 bp, and subsequently cloned into the pTelMV-GW vector. The resulting constructs were named pTelMV-GW-GFP_309_ and pTelMV-GW-GFP_405_, respectively. The 16c plants were agroinoculated with pTelMV-GW-GFP_309_, pTelMV-GW-GFP_405_, or wild-type TelMV (pTelMV) (Gou et al., 2023). As expected, at 18 dpi, the 16c plants inoculated with pTelMV-GW constructs carrying GFP fragments showed apparent red fluorescence with reduced green fluorescence under UV light in the upper leaves compared to the Mock- or pTelMV-infiltrated plants (Figure 4A, upper panel), suggesting the GFP gene is likely silenced in the pTelMV-GW-GFP_309_ and GFP_405_-inoculated 16c plants. This phenomenon is much more obvious at 26 dpi where the systemic leaves of 16c plants inoculated by pTelMV-GW constructs carrying GFP fragments exhibited mainly red fluorescence under UV light (Figure 4A, lower panel), indicating the GFP gene is silenced to a great extent in 16c *N. benthamiana* plants.

**Fig 4.**
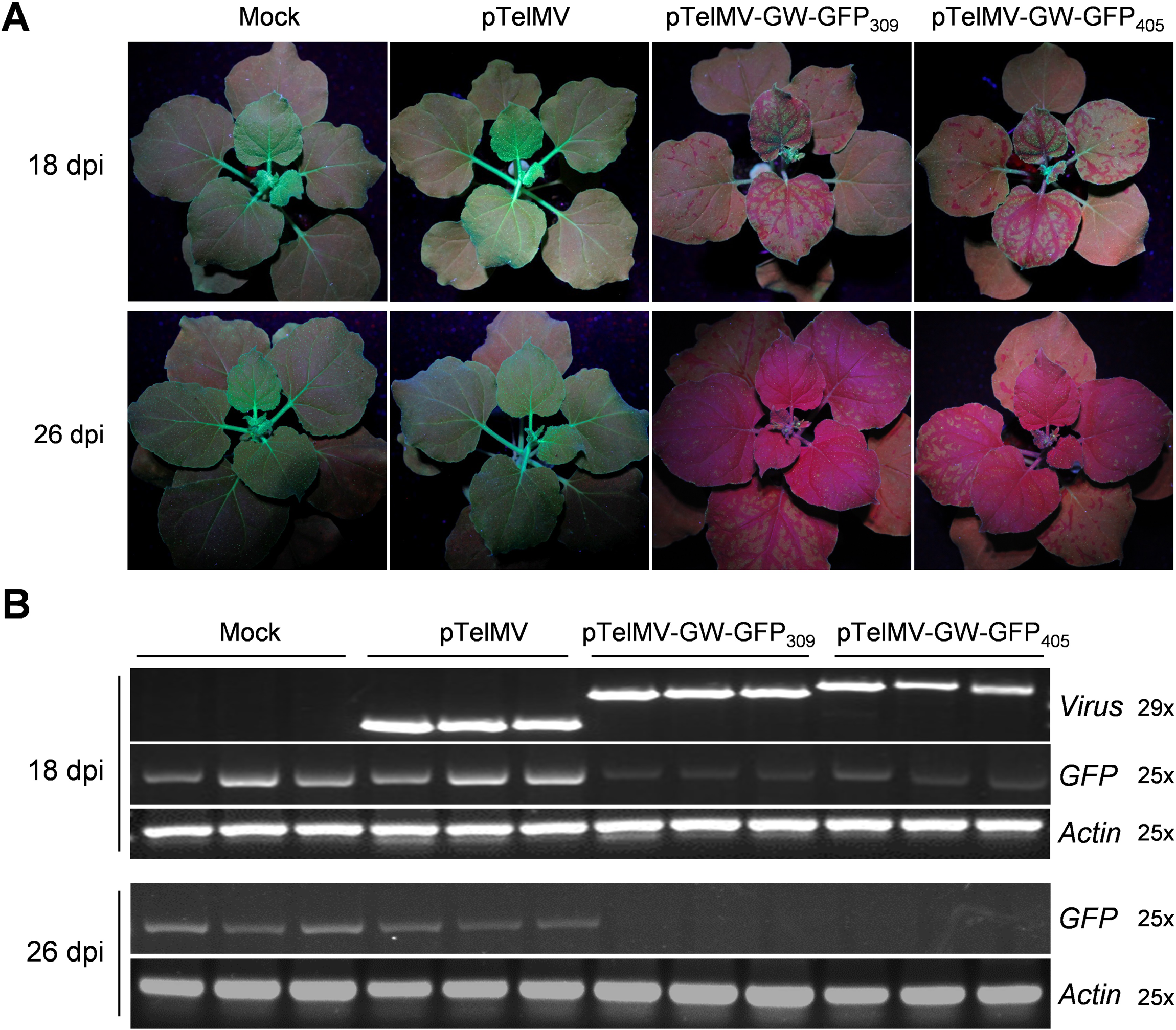
TelMV-based VIGS vector for gene silencing in the GFP-transgenic *N. benthamiana* plants (16c). **A.** 16c plants inoculated by buffer (Mock), wild-type TelMV (pTelMV) or the pTelMV-GW constructs harboring different GFP fragments, under UV light at 18 and 26 dpi, respectively. **B.** RT-PCR analysis of the GFP mRNA level of 16c plants inoculated by various constructs at 18 and 26 dpi, respectively. TelMV- and GFP-specific primers were used to detect viral RNA and GFP mRNA, respectively. The *Actin* gene serves as an internal control.

Next, we extracted total RNA from the upper leaf of each sample and performed RT-PCR analysis. The results revealed a reduced GFP mRNA level of *N. benthamiana* leaf inoculated by pTelMV-GW-GFP_309_ or pTelMV-GW-GFP_405_, compared to that of Mock- or pTelMV-infiltrated plants at 18 dpi (Figure 4B). In addition, the GFP fragments are genetically stable in the systemic leaf evidenced by the amplicons of correct size using the primers flanking the GFP insert that can amplify a single bigger band comprising the full-length GFP insert (Figure 4B). More importantly, at 26 dpi, RT-PCR analysis showed a nearly undetectable mRNA level of GFP (Figure 4B), indicating the GFP gene of 16c plants was silenced upon inoculation of pTelMV-GW constructs carrying GFP fragments. Taken together, we demonstrated that the pTelMV-GW vector is capable of triggering gene silencing in the 16c plants.

### Silencing of *phytoene desaturase* (*PDS*) in passion fruit plants

To test the ability of TelMV to induce endogenous gene silencing in passion fruit plants, we chose the *phytoene desaturase* (*PDS*) gene for the test, as the PDS gene is essential for carotenoid production and its silencing results in photobleaching. A 345 bp fragment of the *PePDS* ORF*, PePDS_345_*, was amplified from the cDNA of passion fruit leaf and cloned into pTelMV-GW, resulting in pTelMV-GW-PDS_345_ (Figure 5A). Passion fruit plants in the first- to second-true-leaf stage were rub-inoculated with sap prepared from pTelMV-GW-PDS_345_-inoculated *N. benthamiana* leaf. For the majority of inoculated passion fruit plants, photobleaching was first observed at about 20 dpr. At 30 dpr, passion fruit plants inoculated with pTelMV-GW-PDS_345_ showed apparent, but not entire photobleaching on the upper, uninoculated leaves (Figure 5B), indicating the endogenous gene *PDS* had been successfully silenced to a certain extent. In contrast, all leaves of plants inoculated with buffer (Mock) or the virus control (pTelMV-GW-GFP_309_) remained green (Figure 5B).

**Fig 5.**
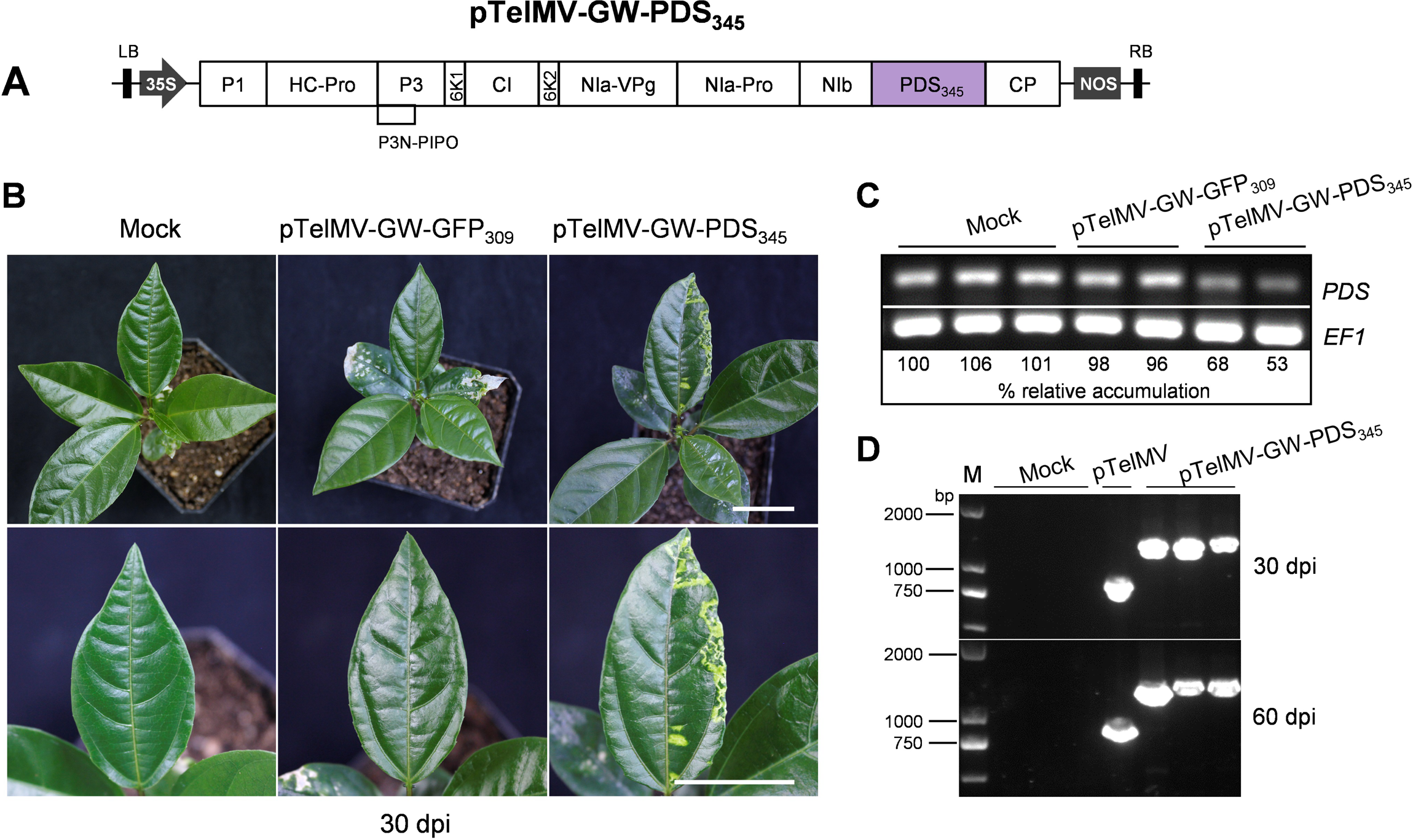
Silencing of endogenous *phytoene desaturase* (*PDS*) gene in passion fruit plants using pTelMV-GW. **A.** Schematic representation of pTelMV-GW-PDS_345_ vector for triggering PDS gene silencing in passion fruit plants. PDS: phytoene desaturase. **B.** Passion fruit plants inoculated by buffer (Mock), pTelMV-GW-GFP_309_ (virus control) or pTelMV-GW-PDS_345_ at 30 dpr. Bars, 2 cm. **C.** RT-PCR analyses of *PDS* mRNA level sampled from passion fruit leaf exhibiting bleaching phenotype. The passion fruit *EF1* gene serves as an internal control. **D.** RT-PCR analysis of RNA samples extracted from pTelMV-GW-PDS_345_-infected passion fruit plants at 30 and 60 dpr, respectively. M: DNA ladder. pTelMV plasmid DNA serves as the wild-type virus control.

Next, the total RNA of passion fruit leaf exhibiting photobleaching was extracted, and RT-PCR was performed to quantify the *PDS* mRNA level using the *EF1* gene as the internal control. The expression level of *PDS* was similar between pTelMV-GW-GFP_309_-infected and Mock-inoculated passion fruit leaves, suggesting TelMV infection alone did not significantly affect *PDS* expression (Figure 5C). In contrast, PDS mRNA was 32-47% less abundant in the bleached leaf sample induced by pTelMV-GW-PDS_345_ infection, than in the Mock-inoculated plants (Figure 5C). Furthermore, the stability of pTelMV-GW-PDS_345_ was investigated by RT-PCR of total RNA samples from passion fruit leaf extracted at 30 and 60 dpi, respectively. The results showed that there were no deletion derivatives (Figure 5D), suggesting that the pTelMV-GW vector retained the inserted PDS sequence upon viral systemic infection in passion fruit plants.

Taken together, the TelMV-based vector, pTelMV-GW successfully triggered the stable silencing of endogenous *PePDS* gene in passion fruit plants, despite the photobleaching was not strong at the early stage.

### Optimization of TelMV-based VIGS Vector by using mild strain

As described above, pTelMV-GW can trigger the endogenous PDS gene silencing in passion fruit plants, but with no strong and decent photobleaching at the early stage. We speculated that this is probably due to the interference of the potyviral encoded HC-Pro protein, a strong RNA silencing suppressor (RSS) (Wang et al., 2024). To test this hypothesis, we improved the pTelMV-GW vector by replacing the wild-type HC-Pro with the point mutated HC-Pro (R to K, at the position of 181) from the previously reported mild strain, TelMV-R181K (Figure 1D). This mild strain encodes a mutant HC-Pro with reduced RSS activity and does not induce noticeable symptoms in plants (Wang et al., 2024). This newly upgraded vector is designated pTelMV-R181K-GW (Figure 1E), a destination vector for the expression of GOI through LR reaction (Figure 1F).

To explore the capability of pTelMV-R181K-GW for endogenous gene silencing in passion fruit plants, two classic VIGS marker gene *PDS* and *ChlI* (*magnesium chelatase subunit I*) fragments (345 bp and 348 bp, respectively) were cloned into this vector, the resulting constructs were named pTelMV-R181K-GW-PDS_345_ and pTelMV-R181K-GW-ChlI_348_. Subsequently, Passion fruit plants were rub-inoculated with sap prepared from these two viral constructs-infected *N. benthamiana* plants to allow for systemic silencing of *PDS* and *ChlI*. As expected, an apparent photobleaching phenotype was observed at an early stage of 20 dpr and, strong and robust photobleaching developed at 30 dpr (Figure 6A). In contrast, Mock and virus control (pTelMV-R181K-GW-GFP_309_)-inoculated passion fruit plants remain green, suggesting the mild strain of TelMV-R181K is capable of triggering efficient endogenous gene silencing in passion fruit plants.

**Fig 6.**
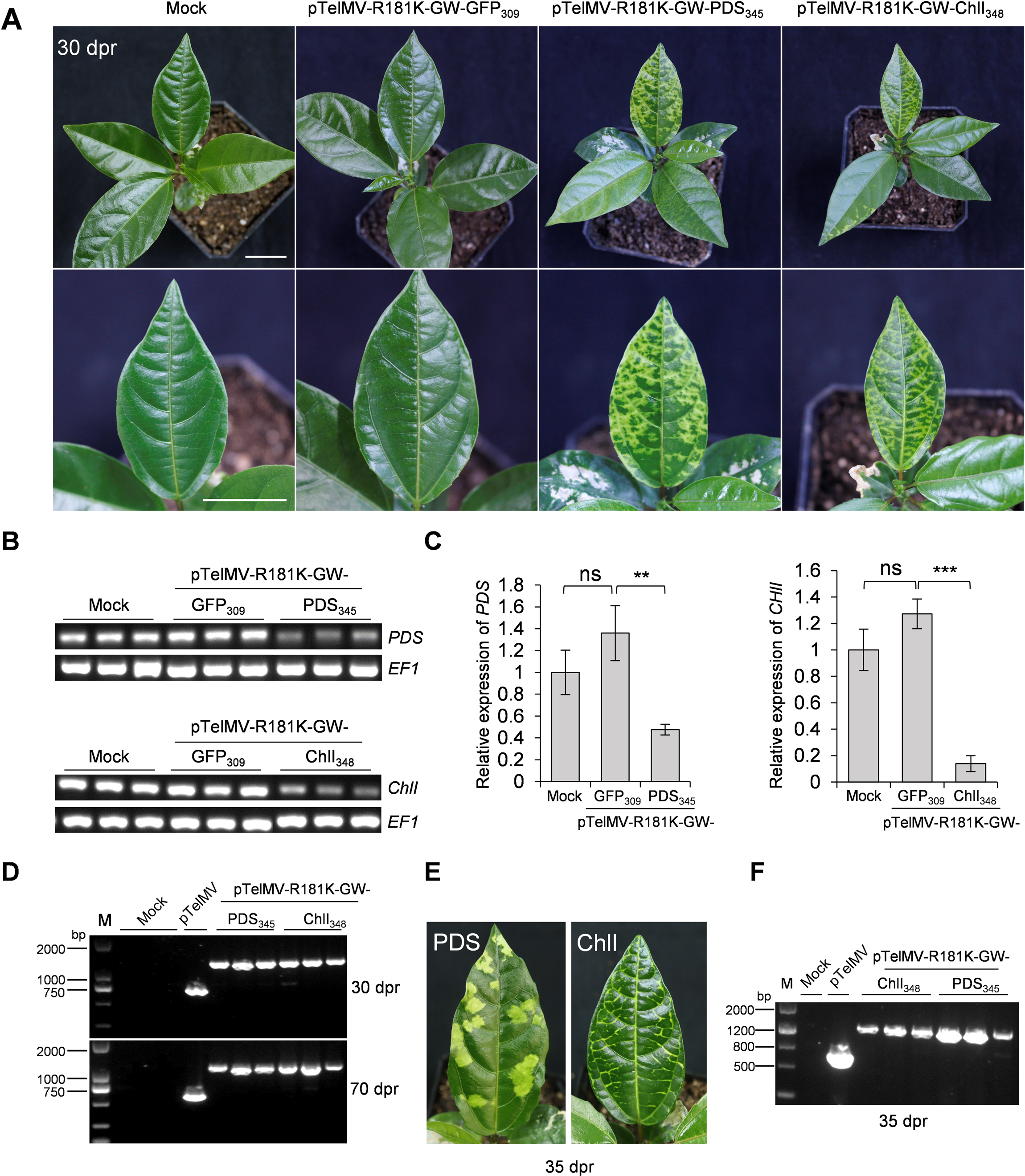
The newly developed pTelMV-R181-GW vector for efficient and stable silencing of endogenous genes in passion fruit plants. **A.** Passion fruit plants inoculated by buffer (Mock), pTelMV-R181K-GW-GFP_309_ (virus control), pTelMV-R181K-GW-PDS_345_ or pTelMV-R181K-GW-ChlI_348_ at 30 dpr. Bars, 2 cm. *ChlI*: *magnesium chelatase subunit I*. **B.** RT-PCR analysis of *PDS* and *ChlI* mRNA level sampled from passion fruit leaf exhibiting bleaching phenotype at 30 dpr. The passion fruit *EF1* gene serves as an internal control. **C.** qRT-PCR analyses of *PDS* and *ChlI* mRNA levels sampled from passion fruit leaf exhibiting bleaching phenotype at 30 dpr. The transcript levels were normalized against the internal control *EF1*. Error bars represent the standard deviation of three biological replicates. Statistically significant differences were determined by an unpaired two-tailed Student’s t test, \*\**p* < 0.01; \*\*\**p* < 0.001; ns: not significant. **D.** RT-PCR analysis of RNA samples extracted from passion fruit plants inoculated by pTelMV-R181K-GW-PDS_345_ or pTelMV-R181K-GW-ChlI_348_ at 30 and 70 dpr, respectively. M: DNA ladder. pTelMV plasmid DNA serves as the wild-type virus control. **E.** Photobleaching and yellowing phenotypes of passion fruit leaves after one serial passage at 35 dpr, using the inoculum of pTelMV-R181K-GW-PDS_345_ or pTelMV-R181K-GW-ChlI_348_, respectively. **F.** RT-PCR analysis of RNA samples extracted from passion fruit plants in the first serial passage of the inoculum of pTelMV-R181K-GW-PDS_345_ or pTelMV-R181K-GW-ChlI_348_. M: DNA ladder. pTelMV plasmid DNA serves as the wild-type virus control.

Next, we extracted the total RNA from the systemic leaf of passion fruit plants and conducted RT-PCR and qRT-PCR analyses. There is no significant difference in *PDS* and *ChlI* mRNA levels between Mock and virus control-inoculated leaves (Figure 6B, C), suggesting virus alone did not significantly affect *PDS* and *ChlI* expression levels in passion fruit plants. However, pTelMV-R181K-GW-PDS_345_ and pTelMV-R181K-GW-ChlI_348_ infection resulted in a significant reduction of *PDS* and *ChlI* expression levels in the photobleached leaves, with an average of 46.8% and 14% of the Mock control plants, respectively (Figure 6B, C).

To explore the stability of *PDS* and *ChlI* inserts during virus multiplication in passion fruit plants, we conducted RT-PCR analysis on total RNA extracted from the systemic leaves exhibiting photobleaching phenotype at 30 and 70 dpr, respectively. A pair of primers flanking the insert was used to detect the intact insert or deletion derivatives.

A single band containing the intact *PDS* or *ChlI* insert can be detected. We did not detect apparent deletion derivatives in a total of 12 individual plants in different periods. These results suggest that the 345-bp *PDS* fragment and 348-bp *ChlI* fragments are stable in the newly developed TelMV-based VIGS vector, pTelMV-R181K-GW.

We further tested whether the silencing effect of pTelMV-R181K-GW-PDS_345_ and pTelMV-R181K-GW-ChlI_348_ could be passaged, and if yes, are the inserts stable during serial passage. The sap from passion fruit leaves displaying obvious photobleaching was used to rub inoculate the healthy passion fruit plants. As expected, photobleaching phenotype could be observed at 12 dpr, and progressively developed at 35 dpr (Figure 6E), demonstrating the *PDS* and *ChlI*-silencing effect could be passaged in passion fruit plants. We also investigated the stability of inserts during serial passage, RT-PCR analysis using primers flanking the insert sites showed a single band containing the intact inserts, suggesting the *PDS* and *ChlI* inserts are stable in passion fruit plants during serial passage (Figure 6F).

Taken together, these data demonstrated that TelMV mild strain (TelMV-R181K)-based vector, pTelMV-R181K-GW can be used as an efficient and stable VIGS vector, and the silencing effect can be serially passaged in passion fruit plants.

## Discussion

This study reports the development of TelMV-based viral vectors for foreign gene expression and endogenous gene silencing in passion fruit plants. First, the pTelMV-GW vector serves as a VOX vector that systemically expresses foreign genes (GFP and CrtB) in the model plant *N. benthamiana*, as well as the tropical fruit-bearing plant, passion fruit (*P. edulis*). In addition, the pTelMV-R181K-GW vector, based on an attenuated TelMV strain, can be applied in endogenous gene silencing in passion fruit, illustrated by efficient systemic gene silencing of *PePDS* and *PeChlI*. To our knowledge, this is the first report on VOX and VIGS vectors in passion fruit plants. The reported TelMV-based vectors provide a valuable tool for gene function analysis in passion fruit plants, without the need for plant transformation.

### TelMV-based vector for transient gene overexpression

The pTelMV-GW VOX vector incorporates the Gateway-compatible recombination sites, thus facilitating the rapid and high-throughput cloning of foreign genes (Fig 1). This study shows that pTelMV-GW can tolerate the foreign sequence insertion at the NIb-CP junction and can carry the 714-nucleotide-long (238 aa) *GFP* gene- and 927-nucleotide-long (309 aa) crtB gene-coding sequence, respectively in both *N. benthamiana* and *P. edulis* plants (Fig 2 and 3). This is consistent with the study by Tuo et al. where the authors reported successful systemic expression of GFP and crtB in cassava using a cassava common mosaic virus (CsCMV, genus *Potexvirus*)-based VOX vector (Tuo et al., 2023). Moreover, we have demonstrated the inserted GFP and crtB fragments are maintained in the TelMV genome during virus replication in passion fruit plants (Fig 2 and 3). This highlights the stability of the foreign insertions while using the pTelMV-GW vector for heterologous protein expression. Stability is a key factor in using potyvirus-based vectors for expressing non-viral proteins. It was reported that deletions of non-viral fragments in this case of a 2147-nucleotide-long GUS (*beta-glucuronidase*) gene, occurred during potyviral replication (Mei et al., 2019).

Potyvirus-based viral vectors have been broadly reported for transient gene overexpression, including TEV, PVA, TuMV, PPV, SCMV, WSMV and ZYMV (Xie et al., 2021). However, potyviral vectors are generally considered to express only moderately sized proteins, ranging from GFP (238 aa), NPT II (264 aa) to GUS (603 aa) (Choi et al., 2000; Bouton et al., 2018; Yang et al., 2024). This is probably due to the relatively large viral genome of potyvirus (around 10 kb) and its polyprotein processing as gene expression strategy. Intriguingly, a recent study reported that a zinnia potyvirus, zinnia mild mottle virus (ZiMMV), successfully expressed a large protein, Cas9 (1368 aa) in the infiltrated leaf but not the upper non-infiltrated leaf (Yang et al., 2024). It is a valid direction for the future to evaluate the carrying capacity of the TelMV-based VOX vector.

### TelMV-based vector for endogenous gene silencing in passion fruit plants

Currently, there is no VIGS tool available for passion fruit plants before our pre-print work and this study (Wang et al., 202). Several reasons might contribute to this, firstly, the chromosome-scale genome assembly of passion fruit plants has been only reported recently (Xia et al., 2021), and the gene function study of passion fruit plants is still in its infancy. Secondly, the most frequently used TRV-based viral vector cannot infect passion fruit plants based on our multiple trials (data not shown). Meanwhile, the infectious clone of passion fruit-infecting virus has been recently constructed, including TelMV, Passiflora mottle virus (PaMoV) and East Asian passiflora virus (EAPV) (Gou et al., 2023; Do et al., 2023; Chong et al., 2023). To date, there are over 30 viruses reported that can naturally infect passion fruit plants, the majority of these viruses belong to the genus of *Potyvirus* (Wang et al., 2024; Zhang et al., 2024).

Here, we have demonstrated that TelMV, a potyvirus, can be developed for an efficient VIGS vector in passion fruit plants. Initially, we showed the potential of TelMV-based vector (pTelMV-GW) as VIGS vector both in *N. benthamiana* and *P. edulis* plants. This is supported by 1) pTelMV-GW containing GFP fragments triggered gene silencing in GFP-transgenic tobacco plants (16c) (Figure 4), and 2) pTelMV-GW expressing a partial fragment of *PePDS*, resulted in photobleached phenotype correlated with the reduced *PDS* mRNA levels, despite of unapparent photobleaching in the early stage in passion fruit seedlings (Figure 5). We then overcome this problem by developing a novel TelMV-based VIGS vector (pTelMV-R181K-GW) by using the attenuated strain TelMV-R181K that harbors HC-Pro point mutation with greatly reduced RSS activity (Wang et al., 2024). The newly developed TelMV VIGS vector efficiently silent *PePDS* and *PeChlI* respectively, in passion fruit seedlings, resulting in photobleached phenotype correlated with the reduced *PDS/ChlI* mRNA levels (only 46.8% and 14% to that of control expression levels, respectively) (Figure 6A, B and C). Furthermore, the insertion is stable in the course of viral replication and systemic movement (Figure 6D) and the silencing effect can be passaged to the new passion fruit seedlings using rub-inoculation through leaf sap prepared from plant inoculated by pTelMV-R181K-GW-PDS_345_ and pTelMV-R181K-GW-ChlI_348_ (Figure 6E).

In line with our study, in 2021, Shen laboratory reported a mild strain of papaya leaf distortion mosaic virus (PLDMV, *Potyvirus*) harboring HC-Pro mutation can be used for VIGS vector for gene silencing in papaya, they successfully silenced five endogenous papaya genes, including, *PDS*, *Mg-chelatase H subunit*, putative *GIBBERELLIN (GA)-INSENSITIVE DWARF1A and 1B* and the cytochrome P450 gene *CYP83B1* (Tuo et al., 2021). Together, the two studies highlight the major role of mild strain harboring HC-Pro mutation with weakened or abolished RNA silencing suppressor (RSS) activity towards potyvirus-based VIGS vectors. Different from this, potyvirus has also been reported for VIGS vector without using HC-Pro mutants, including MDMV, WMV and SCMV (Xie et al., 2021; Houhou et al., 2021; Chung et al., 2022). Taken together, these reports illustrated that potyviruses can be used for VIGS in multiple plant species, including papaya, maize, melon and passion fruit.

The TelMV VIGS vector we developed has several advantages. First, pTelMV-R181K itself can systemically infect passion fruit plants and does not induce visual symptoms that interfere with a VIGS phenotype (Wang et al., 2024). Second, pTelMV-R181K-GW is Gateway-compatible and allows for easy, rapid and high-throughput gene cloning. Lastly, the TelMV VIGS vector offers stable and sustainable gene silencing in passion fruit plants. Nevertheless, more experiments should be carried out to illustrate the capability of the TelMV-based VIGS vector for silencing various passion fruit endogenous genes, especially, genes involved in the biosynthesis of the health-promoting compounds in passion fruit. In addition, as the host range of TelMV has been continuously increasing in recent years (Zhang et al., 2024), TelMV-based VIGS vector is also expected to be applied in other plants, especially for these species lacking simple and efficient functional genetic analysis tools, such as telsoma and patchouli.

## Conclusion

Here, we reported the development of VOX and VIGS vectors (pTelMV-GW and pTelMV-R181K-GW, respectively) in passion fruit plants. The newly-developed TelMV-based vectors are Gateway-compatible, allowing for convenient and rapid molecular cloning, and offer stable heterologous gene expression as well as stable and sustainable silencing of endogenous genes in the perennial woody passion fruit plants. In all, the TelMV viral vector system will provide a powerful and rapid biotechnological tool for discovering horticulturally important genes for improving passion fruit production and quality.

## Materials and Methods

### Plant materials and growth conditions

*N. benthamiana* or *P. edulis* seeds were sown in 7-cm-wide square pots at two-cm depth and germinated in PINDSTRUP substrate supplemented with and vermiculite and perlite (4:1:1). Individual plants were transferred to pots and grown in a growth chamber with 16 h of light at 25°C and 8 h of darkness at 23°C. The relative humidity with set at 65% relative humidity.

### Plasmid construction

pTelMV-GW: The plant expression binary vector pTelMV-GW was based on the previously reported viral vector pPasFru-G (pTelMV-GFP) (Gou et al., 2023). The Gateway recombination frame (Fragment 2, F2) was amplified from pEarleyGate 103 (Invitrogen, CA, USA) using the primer pair c/d (Table 1), and subsequently inserted between the NIb and CP cistron by the following overlapping PCR method: Fragment 1 and 3 (F1, F3) were amplified from pPasFru-G using the primer pairs a/b and e/f (Table 1), respectively. Furthermore, F2 was inserted between the F1 and F3 by the overlapping PCR using primer pair a/f. The resulting fragment was named F4. Lastly, F4 was double digested with *Spe*I and *Avr*II enzymes and subsequently ligated into the predigested backbone vector pPasFru-G to construct the Gateway-based TelMV vector pTelMV-GW.

**Table 1:**
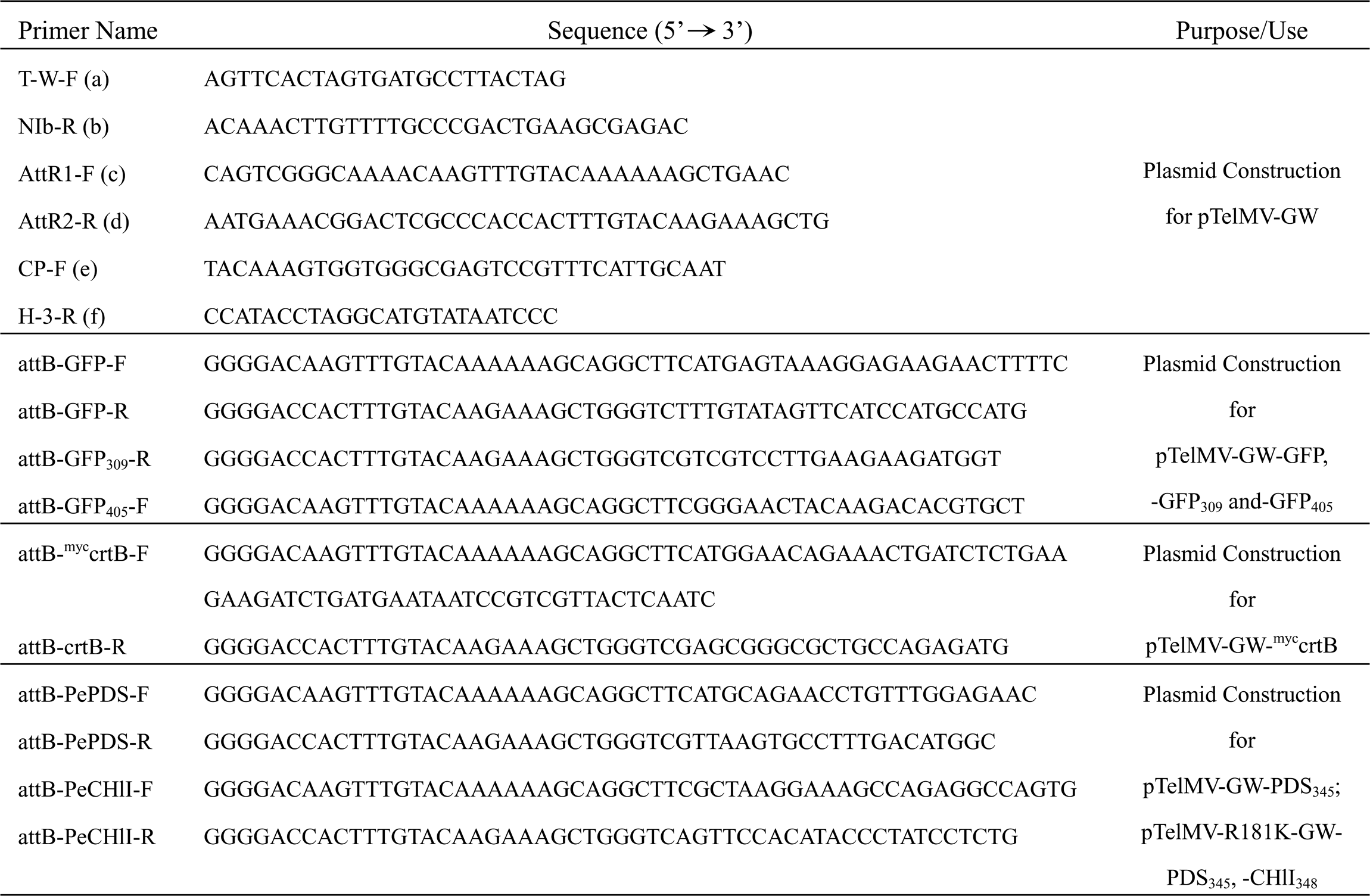
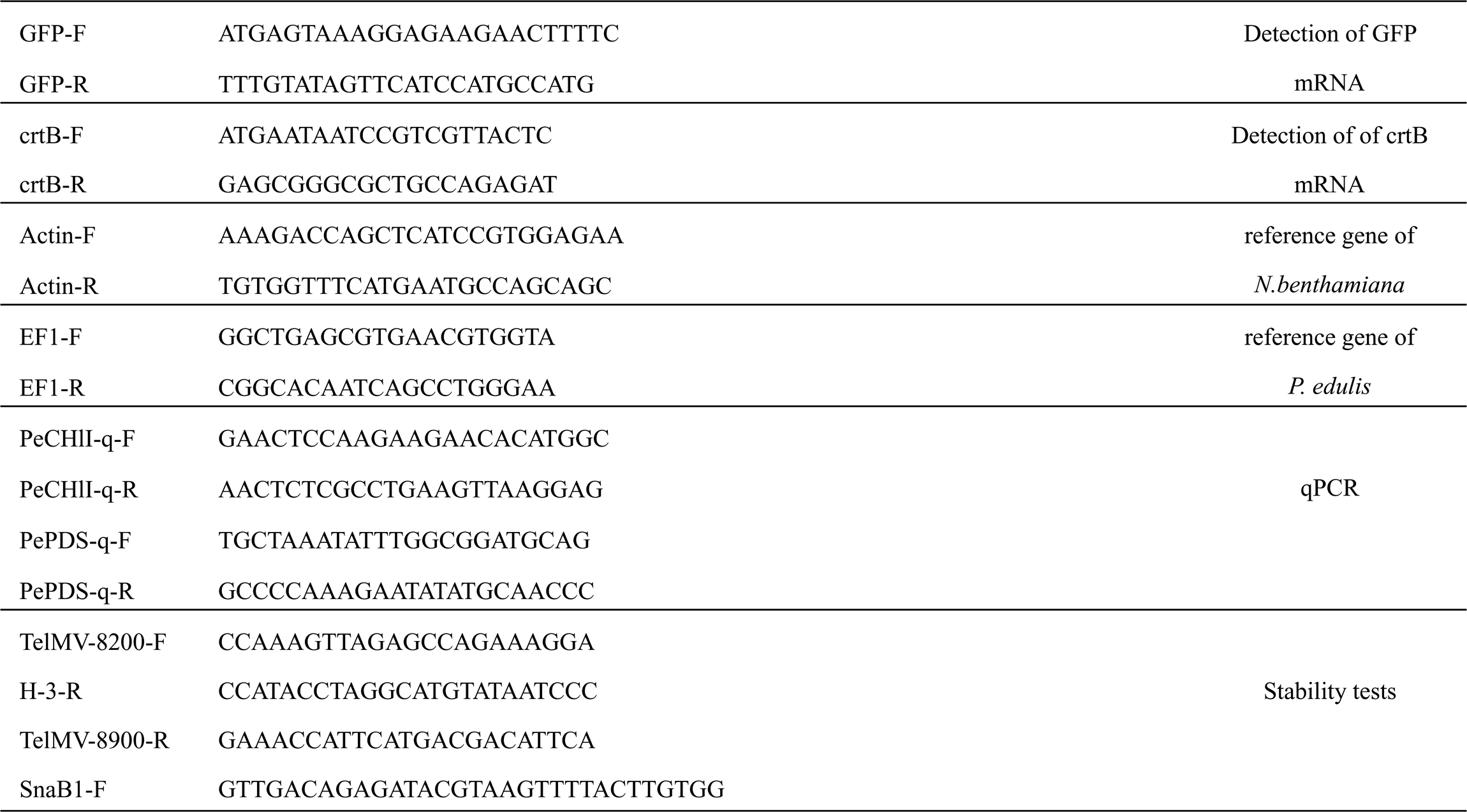
Primers used in this study.

pTelMV-R181K-GW: The backbone of pTelMV-R181K-GW is based on pTelMV-R181K (Wang et al. 2024), which was previously reported by our group in 2024. The Gateway-compatible vector pTelMV-GW was double digested with *Spe*I and *Aat*II enzymes, and subsequently ligated into the predigested backbone vector pTelMV-R181K to obtain pTelMV-R181K-GW.

pTelMV-GW-GFP: The full length of GFP gene was amplified from GFP-tagged TelMV infectious clone (pTelMV-GFP) (Gou et al., 2023) using primer pairs attB-GFP-F and attB-GFP-R (Table 1). The PCR product was then cloned into pDONR221 (Invitrogen, CA, USA) through the BP reaction and subsequently cloned into the destination vector pTelMV-GW by LR reaction.

pTelMV-GW-GFP_309_ and pTelMV-GW-GFP_405_: The full length of GFP was divided into two fragments GFP_309_ and GFP_405_ comprising 309 bp and the remaining 405 bp, respectively for amplification using pTelMV-GFP as the template and primer pairs attB-GFP-F/GFP_309_-R and attB-GFP_405_-F and attB-GFP-R (Table 1). The GFP_309_ and GFP_405_ fragments were recombined into pDONR221 (Invitrogen, CA, USA) using the BP CLONASE enzyme, followed by the LR reaction using pTelMV-GW as the destination vector.

pTelMV-GW-^Myc^CrtB: The complete crtB-coding region was amplified from pCsCMV2-NC (Tuo et al., 2023) by using primer set attB-^myc^crtB-F and attB-crtB-R (Table 1). Note that we added Myc-coding sequence at the N-terminus of the primer attB-^myc^crtB-F. The resulting PCR product was recombined into pDONR221 by BP reaction and subsequently recombined into the destination vector pTelMV-GW by LR reaction.

pTelMV-GW-PDS_345_: As the passion fruit PDS gene (*PePDS*) has not been reported previously, we first did the mining in the genome sequence of *P. edulis* by BLAST searching using either tobacco PDS gene (*NbPDS*) or Arabidopsis PDS gene (*AtPDS*) as the query. Next, multiple sequence alignments were performed among the putative *PePDS, NbPDS* and *AtPDS*. The primer set attB-PePDS-F/attB-PePDS-R (Table 1) for amplifying *PePDS* fragment (345 bp) was designed from the conserved nucleotide sequence of *PePDS*. The PCR product was further used for cloning to the destination vector pTelMV-GW through classic BP and LR reactions.

pTelMV-R181K-GW-GFP_309_, -PDS_345_ and -ChlI_348_: To construct pTelMV-R181K-GW-GFP_309_ and PDS_345_, the pDONR221-GFP_309_ and PDS_345_ described above are recombined into the destination vector pTelMV-R181K-GW through LR reaction. Moreover, a 348 bp-fragment of ChlI gene was amplified from the cDNA of passion fruit plant by using the primer set attB-PeCHlI-F/attB-PeCHlI-R (Table 1). The resulting PCR product was subsequently cloned into the destination vector pTelMV-R181K-GW through BP and LR reactions.

### Plant Inoculation

For the inoculation of *N. benthamiana* plants, *Agrobacterium tumefaciens* harboring virus infection clones were used for leaf infiltration. In brief, *A. tumefaciens* strain GV3101 harboring various plasmids was grown overnight on a shaker at 200 rpm and the bacterial culture was then centrifuged, washed and finally resuspended in the agroinfiltration buffer (10 mM MgCl_2_, 10 mM MES, pH 5.7) supplemented with 200 μM acetosyringone. The optical density (OD600) of the bacterial suspension was then adjusted to 0.5 for infiltration using a needle-less syringe.

For the inoculation of *P. edulis* plants, we used the rubbing method. More specifically, the sap prepared from *N. benthamiana* plants infiltrated with agrobacterium harboring relevant plasmid was rub-inoculated on the cotyledons or true leaves of *P. edulis* plants.

### RNA extraction, RT-PCR and qRT-PCR

Total RNA was extracted from 30 mg leaf tissue of *N. benthamiana* or *P. edulis* plants using TRNzol universal reagents (Tiangen). For first-strand cDNA synthesis, 500 ng of RNA was treated by DNase I (Thermo Fisher Scientific), followed by reverse transcription reaction using the SuperScript III First-strand Synthesis System (Thermo Fisher Scientific) as instructed.

For RT-PCR analyses, primers pairs GFP-F/GFP-R and crtB-F/crtB-R (Table 1) were used to detect the GFP and crtB mRNA (Fig 2C and Fig 3C), respectively. A fragment of *EF1* was amplified from passion fruit cDNA using primers EF1-F/EF1-R (Table 1) and served as the internal control. Primers Actin-F and Actin-R (Table 1) were used for amplifying a fragment of *N. benthamiana Actin* gene and served as the internal control. Primers TelMV-8200-F and TelMV-8900-R (Table 1), located on NIb- and CP-coding regions, respectively, were used to detect viral mRNA or its derivatives for Fig 4B. For the insertion stability test, primer pairs TelMV-8200-F/TelMV-8900-R, SnaBI-F/TelMV-8900-R and SnaBI-F/H-3-R (Table 1) were used for Fig 2F/6D, Fig 3D and Fig 6F, respectively.

For qRT-PCR analysis, qTOWER3 real-time PCR thermal cycler (Analytic Jena AG) with SuperReal PreMix Plus (SYBR green) (Tiangen) was used following the manufacturer’s instructions. Primer pairs PePDS-q-F/PePDS-q-R and PeCHlI-q-F/PeCHlI-q-R are used for the detection of the PDS and ChlI genes in passion fruit plants. The *EF1* transcripts were used as an internal control to normalize the data.

### Western blot analysis

The leaf tissue of *N. benthamiana* or *P. edulis* plants was ground in the protein extraction buffer (50 mM Tris–HCl (pH 6.8), 4% (w/v) SDS, 50 mM dithiothreitol, 10% (v/v) glycerol, 1% (w/v) polyvinylpyrrolidone 40, and 5% (v/v) phenylmethylsulfonyl fluoride) using a pestle and mortar. Subsequently, debris was removed by centrifugation and the supernatant was subjected to immunoblot analysis after SDS-PAGE. Briefly, protein samples were separated on standard SDS-PAGE (12% acrylamide) gel and subsequently transferred to PVDF membrane (Bio-Rad) using the Trans-Blot SD Semi-Dry Transfer Cell (Bio-Rad). Immunoblotting was conducted with anti-GFP or anti-Myc polyclonal antibody (Abcam) as the primary antibodies. The horseradish peroxidase (HRP)-conjugated goat anti-rabbit antibody (Abcam) served as the secondary antibody. Lastly, the blotted PVDF membranes were washed in PBST three times and visualized with HRP substrate using a chemiluminescence system as instructed (Thermo Fisher Scientific).

## Data Availability Statement

The original contributions presented in the study are included in the article, further inquiries can be directed to the corresponding author/s.

## Acknowledgments

We thank Dr. Aiming Wang (Agriculture and Agri-Food Canada) for critical reading of the manuscript, Dr. Runmao Lin (Hainan University) for helping the mining in the genome sequence of *P. edulis* for *PDS* gene and Dr. Justice Norvienyeku (Hainan University) for helpful discussion.

## Author Contributions

ZD, HC, XW and LQ designed the experiment and wrote the manuscript. XW and LQ performed the experiments. All authors analyzed, discussed the data, read, and approved the final manuscript.

